# Sex Identification in Cattle, based on Amelogenin Gene

**DOI:** 10.1101/2021.07.11.451984

**Authors:** Aftab Ahmad, Muhammad Israr, Murad Ali Rahat, Adnan Wahab, Subhan Uddin, Akhtar Rasool, Fazal Akbar, Muzafar Shah

**Affiliations:** Centre for Animal Sciences and Fisheries, University of Swat, Charbagh, Pakistan; Department of Genetics, Hazara University Mansehra, Pakistan; Department of Zoology, Islamia College Peshawar, Pakistan; Centre for Biotechnology and Microbiology, University of Swat, Charbagh, Pakistan; Department of Forensic Sciences, University of Swat, Charbagh, Pakistan

**Keywords:** Cattle blood, DNA Extraction, PCR assay, *Amelogenin* gene, Sex identification

## Abstract

Sex identification is considered an important step in the field of forensic sciences, wildlife and livestock breeding management. In the current experiment we used *Amelogenin* gene as a biological marker for polymerase chain reaction test to identify the sex of cattle from blood remnants, collected at slaughter house. Due to the conserved region of the gene on both sex chromosomes (X and Y) a single primers pair was employed to amplify the gene in a single polymerase chain reaction. In case of band patterns, a 178 base pair fragment for *AMELY* and a 241 base pair fragment for *AMELX* genes were produced. The primer’s competence and exactness was initially checked on known gender cattle samples and then applied to unknown cattle samples for the validation of the experiment. PCR amplicons of unknown gender showed only one band (241-bp) for female DNA and two bands (241-bp, 178-bp) for male DNA, on the platform of agarose gel upon electrophoresis. Our findings showed that the PCR protocol based on *AMELX/Y* gene is a reliable technique for the identification of cattle sex.

## Introduction

Sex identification or slaughter discrimination of cattle has always been prioritized by public in the context of processed food products utilization. The economic aspects of such policy also accept and designate that female cattle meat is less standard than male cattle meat and is therefore yield lower prices in the market (Zeleny and Schimmel, 2002). This is highly prioritized in India, the cow slaughtering is prohibited for meat consumption because of their laws and beliefs in religion. This is quite necessary to put labeling on meat products for the reliability of consumers. Before implementing regulations regarding such issues, the analysts needs to provide a method which must be simple and reliable for the identification of meat products, or to trace back a forensic sample to its origin. A wide range of methodologies, based on PCR has been established for the correct identification of the mammals gender e.g. ZFX/ZFY zinc-finger region (Fredsted and Villessen, 2004), SRY-locus (Bryja and Konecny, 2003). The BRY-1 (bovine specific repetitive sequences) (Matthews and Reed, 1992), BOV97M (Single Copy Sequence) by (Miller and Koopman, 1990) and amelogenin gene (Pfeiffer and Brenig, 2005; Villesen and Fredsted, 2006). Other studies, like TaqMan (Parati et al ., 2006) and CYBR Green Ballin and Madsen, 2007) have also been reported in the innovative studies for sex determination. However the technique based on the amelogenin system has gained greater attention due to fast, reliable, sensitive and accurate outcomes (Tagliavini et al., 1993).

The amelogenin gene code for proteins known as matrix protein (enameline), found in all mammalian tooth enamel. Enamel appears to be the outer, hard and white protective covering of each tooth (Sedwick, 2009). It is used for sex identification because of homologous sequences of the gene are present upon both (X and Y) sex linked-chromosomes (Nakahori et al., 1991) and unlike other techniques this one only require a set of specific primers to a particular species for identification. Although the gene is amplified in a single polymerase chain reaction, in which a set of forward and reverse primers specific to cattle has been designed to produce amplicons of both the conserved regions in a single thermal cycler reaction for achieving different PCR results (Roffey et al., 2000). Two fragments 280 and 217-bp have shown in a study conducted by (Ennis and Gallagher, 1994) and (Chen et al., 1999) amplified 417 and 340-bp. In this experiment we expanded the work over sex identification of cattle through the *amelogenin* gene while isolating the genetic material from blood remnant at slaughter house, which is novel to the same line of work accomplished previously for sex identification.

## Materials and Methods

### 2.1. Collection of samples

Whole blood samples were obtained from 18 cattle. Of which two were taken as a referenced (male and female cattle samples) and the rest of the samples were collected randomly at a local slaughter house of District Swat. The blood samples were collected immediately in 3ml EDTA tubes after the slaughtering of the cattle and processed for storage at 4°C temperature for further analysis.

### 2.2. Genetic materials isolation

Genetic materials were isolated from 200 μl blood (whole) using DNA extraction kit, (*WizPrep™ gDNA Mini Kit (blood)*, Cat# W71050-100), wizbiosolutions, South Korea. Instructions were followed as mentioned by the manufacturer. The DNA elution was carried out in 50 μl Tris EDTA buffer before visualizing upon electrophoresis over a 0.8% gel (agarose) stained via ethidium bromide and documented through gel documentation system using OmniDoc software (Version 1.1.3.9) as shown in figure 1 and then stored at −20°C for further experiments.

**Figure 1:**
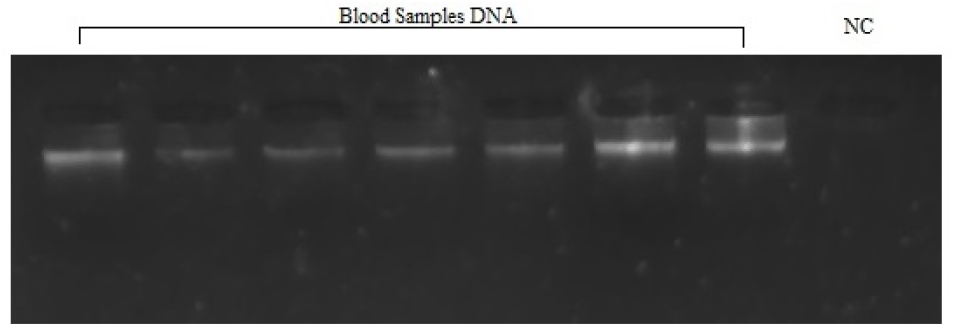
Genomic DNA on 0.8%agarose gel, extracted from cattle blood tissues, whereas NC stands for negative control.

### 2.3. PCR amplification

A pair of primer reported by (Gokulakrishnan et al., 2012), AMEL-F 5′-GGCCAACACTCCATGACTCCA-3′ and AMEL-R 5′-TGGGGAATATYGGAGGCAGAG -3′ was aimed to successfully produce amplicons of the amelogenin gene homologs present on X-chromosome (241-bp) and Y-chromosome (178-bp) in the thermal cycler (PERKIN ELMER, USA). The test/s was carried out in 10 μl mixtures containing each primer of volume 0.25 μl, 2x PCR Taq polymerase (Wiz Bio Solutions) of volume 5.0 μl, 3.5 μl water free of nucleases and 1.0 μl DNA template. Initial denaturation of PCR system was set for 4 min @ 94°C, 34 cycles of denaturation, annealing and extension were performed for 30 sec @ 94°C, 59°C and 72°C respectively. Final extension of 5 min @ 72°C and final hold of the thermal cycler was set for infinity @ 25°C. The PCR amplicons were then electrophoresed on the platform of 2.0% agarose gel.

## Results

The DNA was successfully isolated from the blood remnant samples following manufacturer instructions and visualized under UV light as shown as in figure 1. After PCR amplification, two types of (241 bp, 178 bp) bands were observed in male individuals while female having a single band of 241 bp (Figure 2). The amplicons were observed via 2% agarose gel in which ethidium bromide was used as staining agent. A 50 bp DNA marker (GENEONE Cat# 300009) was simultaneously electrophoresed to assess the amplicons size. The agarose gel was then visualized in the trans-illumination system and the size of the amplicons were determined and documented through OmniDoc software (Version 1.1.3.9).

**Figure 2:**
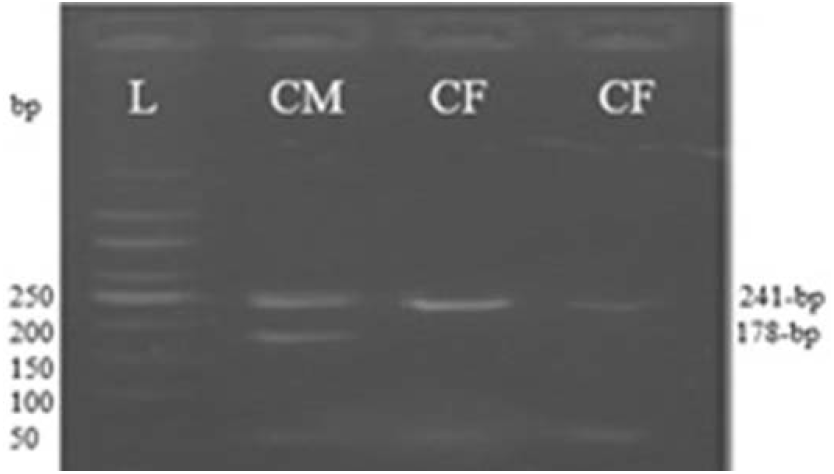
PCR amplicons on 2% agarose gel, L stands for 50bp DNA Marker, CM-for cattle male and CF for cattle female samples, amplicons.

## Discussion

*Amelogenin* gene code for enameline proteins, whom concern is with tooth development and belongs to extra cellular matrix proteins family (Gibson, 1999). The discriminative character of the amelogenin gene is the presence of AMELX and AMELY markers on both sex linked chromosomes (Gokulakrishnan et al., 2012) with a variation of 63 bp sequence in AMEL found over Y chromosome (due to deletion at exon number 5). Hence producing two bands of different sizes for male and a single band for female. This variation may be due to an insertion at that specific point over X-linked chromosome (AMELX) due to which the generated amplicons are of larger size than Y-linked chromosomes. Because of the bands variation between AMELY and AMELX, the amelogenin gene is employed as a biological marker to the purpose of sex identification in almost all mammals such as human (Iwase et al., 2003) cattle (Gibbon et al., 2009), horse (Weikard and Kuhn, 2006) sheep and goat (Pfeiffer and Brenig, 2005).

Specie-specific Cattle-AMEL primers were employed to produce amplicons for both alleles in a single PCR test showing results for both male and female cattle samples on 2% agarose gel through electrophoresis. The primers employed in this study was selected from the study of (Gokulakrishnan et al., 2012). Whom accuracy was evaluated and verified by the assay of extracting genomic DNA from two referenced (male and female) and sixteen random blood samples. The PCR products obtained by this assay showed 178 bp band for Y chromosome and 241 bp band for X chromosome which determined the male and female cattle samples accurately. Our experiment results are also similar to, reported by (Gokulakrishnan et al., 2012). Some different results are also found from previous studies regarding the same gene, amelogenin. But different primers were used which amplified 280bp for AMELX and 217 bp for AMELY alleles (Ennis and Gallagher, 1994). Chen et al (1999) suggested primers which amplified product of 417 bp for AMELX and 340 bp for AMELY chromosomes in cattle. Our findings are similar to the study of (Gokulakrishnan et al., 2012) because of similar primers sequences was used and discrepant with (Ennis and Gallagher, 1994; Chen et al., 1999) because of different primers sequences for the same gene were selected in their studies.

## Conclusions

In the conclusions of our research experiment, it was found that the assay based on PCR of the *amelogenin* gene was found to be a reliable method for identification of sex in cattle from blood. Because, the short amplicons size of the gene (<250-bp) can be compared to earlier work and the approach can be highly applicable in terms of studying degraded DNA. Zero ambiguity found in the assay because the PCR amplicons of both AMELX and AMELY have a good difference between them which was prominent on agarose gel, ensuring the reliability. Feasibility of the study is no additional primers for somatic DNA marker is required and nor post-PCR processes like RFLP is required for the sex identification. This approach of sex identification from cattle blood was proved suitable for routine tests because of its feasibility, cheapness and reliability.

